# Uncoupled activation and cyclisation in catmint reductive terpenoid biosynthesis

**DOI:** 10.1101/391953

**Authors:** Benjamin R Lichman, Mohamed O Kamileen, Gabriel R Titchiner, Gerhard Saalbach, Clare E M Stevenson, David M Lawson, Sarah E O’Connor

## Abstract

Terpene synthases typically form complex molecular scaffolds by concerted activation and cyclization of linear starting materials in a single enzyme active site. Here we show that iridoid synthase, an atypical reductive terpene synthase, catalyses the activation of its substrate 8-oxogeranial into a reactive enol intermediate but does not catalyse the subsequent cyclisation into nepetalactol. This discovery led us to identify a class of nepetalactol-related short-chain dehydrogenase enzymes (NEPS) from catmint (*Nepeta mussinii*) which catalyse the stereoselective cyclisation of the enol intermediate into nepetalactol isomers. Subsequent oxidation of nepetalactols by NEPS1 provides nepetalactones, metabolites that are well known for both insect-repellent activity and euphoric effect in cats. Structural characterisation of the NEPS3 cyclase reveals it binds to NAD^+^ yet does not utilise it chemically for a non-oxidoreductive formal [4+2] cyclisation. These discoveries will complement metabolic reconstructions of iridoid and monoterpene indole alkaloid biosynthesis.

## Main paper

Nepetalactones **1** are volatile natural products produced by plants of the genus *Nepeta*, notably catmint (*Nepeta mussinii* syn *racemosa*) and catnip (*N. cataria*) (Fig. 1a)^1,2^. These compounds are responsible for the stimulatory effects these plants have on cats^3–5^. Moreover, certain insects use nepetalactones as sex pheromones, so production of these compounds by the plant also impacts interactions with insects^6^. Notably, the bridgehead stereocentres (carbons 4a and 7a) vary between^2^ and within^2,4,7^ *Nepeta* species. *N. mussinii* individuals, for example, produce different ratios of cis-trans **1a**, cis-cis **1b** and trans-cis-nepetalactone **1c**^7^. Variation in stereoisomer ratio may influence the repellence of insect herbivores^8,9^. While the ratio of stereoisomers may be responsible for important biological effects, the mechanism of stereocontrol in nepetalactone biosynthesis is not known.

**Figure 1:**
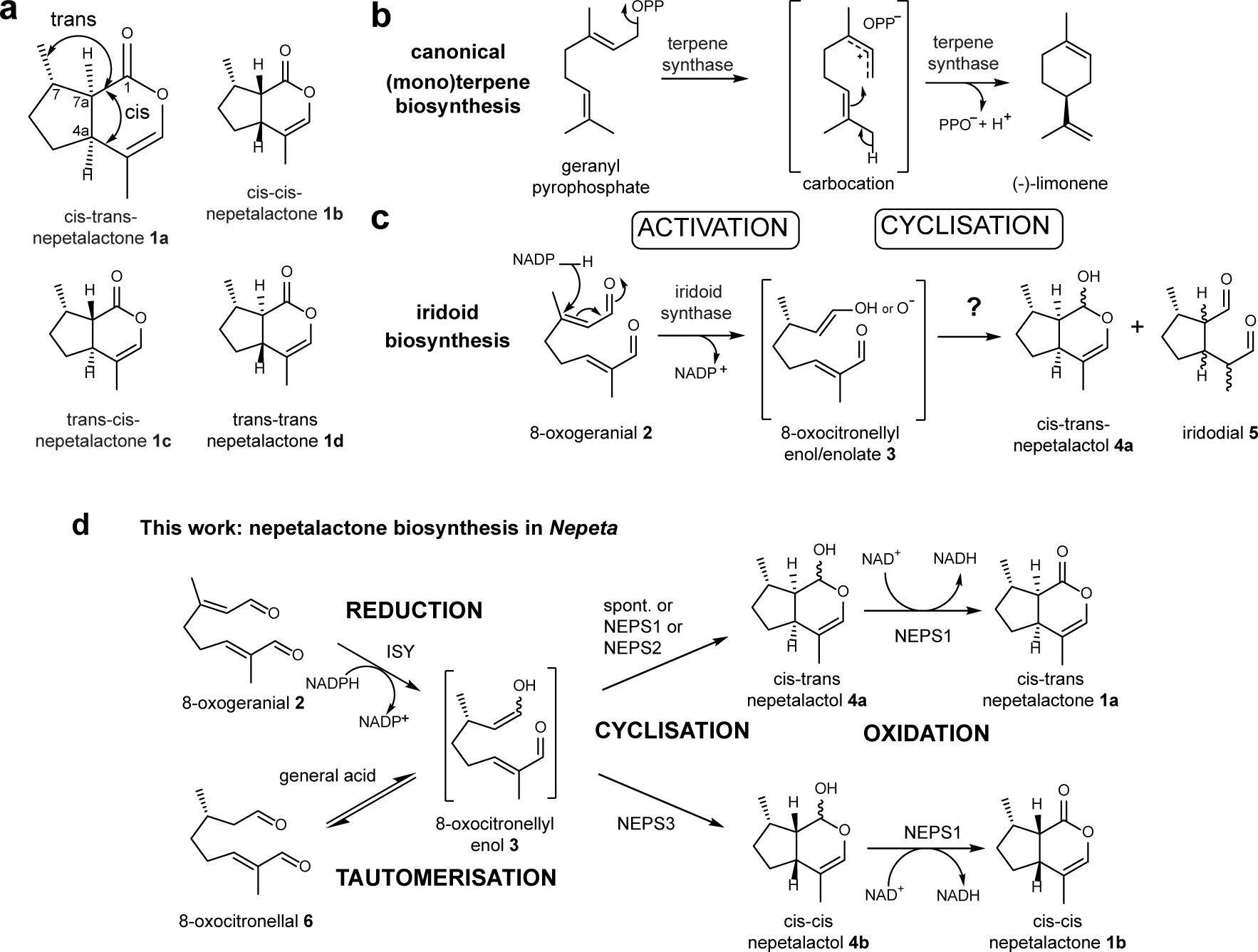
Nepetalactones and terpenoid biosynthesis. **a**, Nepetalactone **1** stereoisomers observed in *Nepeta* species. **b**, Representative canonical terpene biosynthesis mechanism. Typical terpene synthases activate linear precursors by removal of diphosphate or protonation. The activated carbocation intermediates undergo selective cyclisation inside the same terpene synthase active site. Limonene synthase is depicted as an example. **c**, Iridoid biosynthesis mechanism. Iridoid synthase activates its linear precursor (8-oxogeranial **2**) by reduction to form the 8-oxocitronellyl enol/enolate intermediate **3**. This then undergoes cyclisation to form a mixture of cis-trans-nepetalactol **4a** and iridodials **5**. **d**, Biosynthetic origin of cis-trans and cis-cis-nepetalactone stereoisomers in *Nepeta* as reported in this paper.

Nepetalactones are iridoids, non-canonical monoterpenoids containing a cyclopentanopyran ring. Canonical cyclic terpenoids (e.g. (-)-limonene) are biosynthesised from linear precursors by terpene synthases (Fig. 1b)^10^. These enzymes activate linear precursors either by loss of pyrophosphate or protonation^10–12^. The resulting carbocations generated cyclise rapidly to form an array of cyclic products^13^. Therefore, in canonical terpenoid biosynthesis, activation and cyclisation of precursors are coupled and occur in the same enzyme active site.

In plant iridoid biosynthesis, geranyl pyrophosphate is hydrolysed and oxidised into 8-oxogeranial **2**^14^. This precursor then undergoes a two-step activation-cyclisation process, analogous to canonical terpene synthesis (Fig. 1c)^15^. Unlike canonical terpene synthesis, however, activation is achieved by reduction, and the intermediate is not a carbocation, but the enol or enolate species **3**. Cyclisation of this intermediate yields cis-trans-nepetalactol **4a** along with iridodial side products **5** (Fig. 1c).

The conversion of **2** to **4a** and **5** is catalysed by iridoid synthase (ISY)^15^. ISY was first discovered in *Catharanthus roseus* (*Cr*ISY) where it forms part of the biosynthetic route to the anti-cancer monoterpene indole alkaloids vincristine and vinblastine^15,16^. Subsequent studies revealed ISYs from Olive (*Olea europaea*, *Oe*ISY)^17^ and Snapdragon (*Antirrhinum majus*, *Am*ISY)^18^, as well as paralogous proteins in *C*. *roseus* with promiscuous ISY activity^19^. Recently, we identified ISYs from *Nepeta*^20^.

The enzymatic control of the initial reductive activation step has been structurally characterised in *Cr*ISY: crystal structures with cofactor and inhibitor or substrate show binding modes conducive to reduction and formation of an enolate intermediate^21–23^. Furthermore, this reduction is stereoselective, as exemplified by the comparison of *Cr*ISY, which produces 7*S*-**4a**, with *Am*ISY, which produces the enantiomer^18^. In contrast, it remains ambiguous how the cyclisation step controls the stereochemistry of the bridgehead 4a-7a-carbons of the iridoid scaffold to generate the stereochemical variation observed in *Nepeta*.

We have now determined the biosynthetic route to two nepetalactone stereoisomers in *N. mussinii*, cis-trans **1a** and cis-cis **1b**. The discovery of these genes reveals that the reduction and cyclisation steps of iridoid biosynthesis in *Nepeta* are uncoupled and catalysed by distinct enzymes (Fig. 1d). This process involves the diffusion of the activated intermediate 8-oxocitronellyl enol **3** between enzyme active sites. We have discovered and characterised three cyclases (NEPS1-3) from *N. mussinii* that are responsible for the stereoselective cyclisation and subsequent oxidation of activated intermediate **3** into distinct nepetalactone diastereomers. We have also determined the crystal structure NEPS3, providing insight into its mechanism and evolution from a reductase into a redox-neutral cyclase.

## Results

### The mechanism of iridoid synthase (ISY)

Recently we hypothesised that synthesis of the different nepetalactone stereoisomers in *Nepeta* was controlled by species-specific ISYs catalysing both the reduction of **2** and the subsequent stereodivergent cyclisations. However, this was not the case; *Nepeta* ISYs produced the same stereoisomeric product profile as *Cr*ISY^20^. Therefore, an alternative mechanism for the control of iridoid stereochemistry was developed.

The first step toward understanding the origin of the divergent stereochemistry of nepetalactones **1** was to further explore the ISY mechanism. As observed in previous studies^15,18–20^, the ISY catalysed reduction of 8-oxogeranial **2** generates a number of isomeric products (Fig. 2a and b, Supplementary Fig. 1). The product profile of the observed isomers is largely independent of the ISY employed, despite modest sequence identities (48-65%) and different stereoselectivities of the reduction step (*Am*ISY is 7*R*-selective, *Nm*ISY2 and *Cr*ISY are 7*S*-selective) (Fig. 2b).

**Figure 2:**
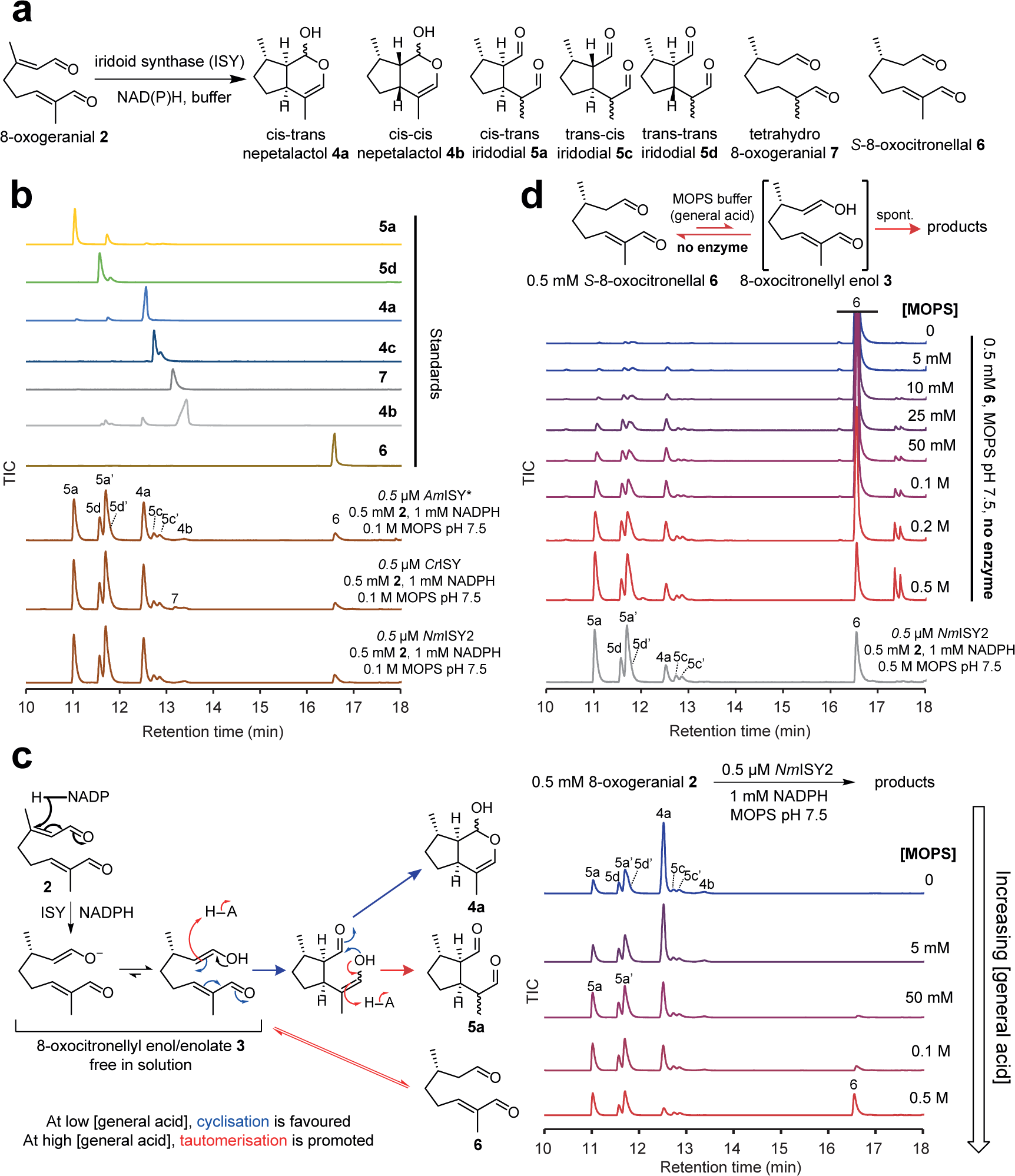
Iridoid synthase (ISY) reaction mechanism. **a**, Product mixture observed in the ISY catalysed reductive cyclisation of 8-oxogeranial **2**. **b**, 8-oxogeranial **2** reduction catalysed by ISYs from *Antirrhinum majus* (*Am*ISY), *Catharanthus roseus* (*Cr*ISY) and *N. mussinii* (*Nm*ISY2). The enzymes form nearly identical product mixtures despite their evolutionary distance and sequence identity. \**Am*ISY produces the opposite enantiomeric series to that depicted in panel **a**^18^. Reactions were incubated for 3 h. See Supplementary Fig 1 for electron ionisation (EI) spectra for compound identification. **c**, ISY catalysed reduction of **2** at different buffer concentrations. At low buffer concentrations, the main product is cis-trans-nepetalactol **4a**. As buffer concentrations increase, higher quantities of cis-trans-iridodials **5a** are observed. At high MOPS (buffer) concentrations (>100 mM) 8-oxocitronellal (**6**) becomes a major product. We propose ISY reduces **2** and then releases the activated **3** into the solvent, where cyclisation occurs. Buffer appears to act as a general acid catalyst, promoting tautomerisation in place of cyclisation. Reactions were incubated for 3 h. See Supplementary Figs 2, 3 and 4 for further exploration of the solvent conditions. **d**, General acid catalysed cyclisation of *S*-8-oxocitronellal **6**. Non-enzymatic formation of **4a** and **5** achieved via tautomerisation and spontaneous cyclisation catalysed by general acid (buffer). The product profile at 0.5 M buffer mimics the equivalent ISY catalysed reduction, supporting a non-enzyme catalysed cyclisation. Reactions were incubated for 16 h. See Supplementary Fig 5 for reactions with **6** in acidic/alkaline unbuffered water and Supplementary Fig 6 for EI spectra. All reactions are presented as GC-MS total ion chromatograms (TICs).

Although the product profile was invariant with respect to the enzyme catalyst, it was strongly influenced by the reaction conditions, especially the concentration of the buffer (Fig. 2c). With low buffer concentration, cis-trans-nepetalactol **4a** was the major product; as buffer concentration was increased, more iridodials **5** and non-cyclised 8-oxocitronellal **6** were observed. A similar pattern was observed with all ISYs and buffers tested (Supplementary Figs. 2a-d, 3). The trend was not a result of cis-trans-nepetalactol **4a** ring opening in high buffer concentrations (Supplementary Fig. 4a)^24,25^. Furthermore, changing the buffer pH also affected the product profile, with lower pH resulting in the formation of more cis-trans-iridodials **5a** compared to cis-trans-nepetalactol **4a** (Supplementary Fig. 2e).

**Figure 3:**
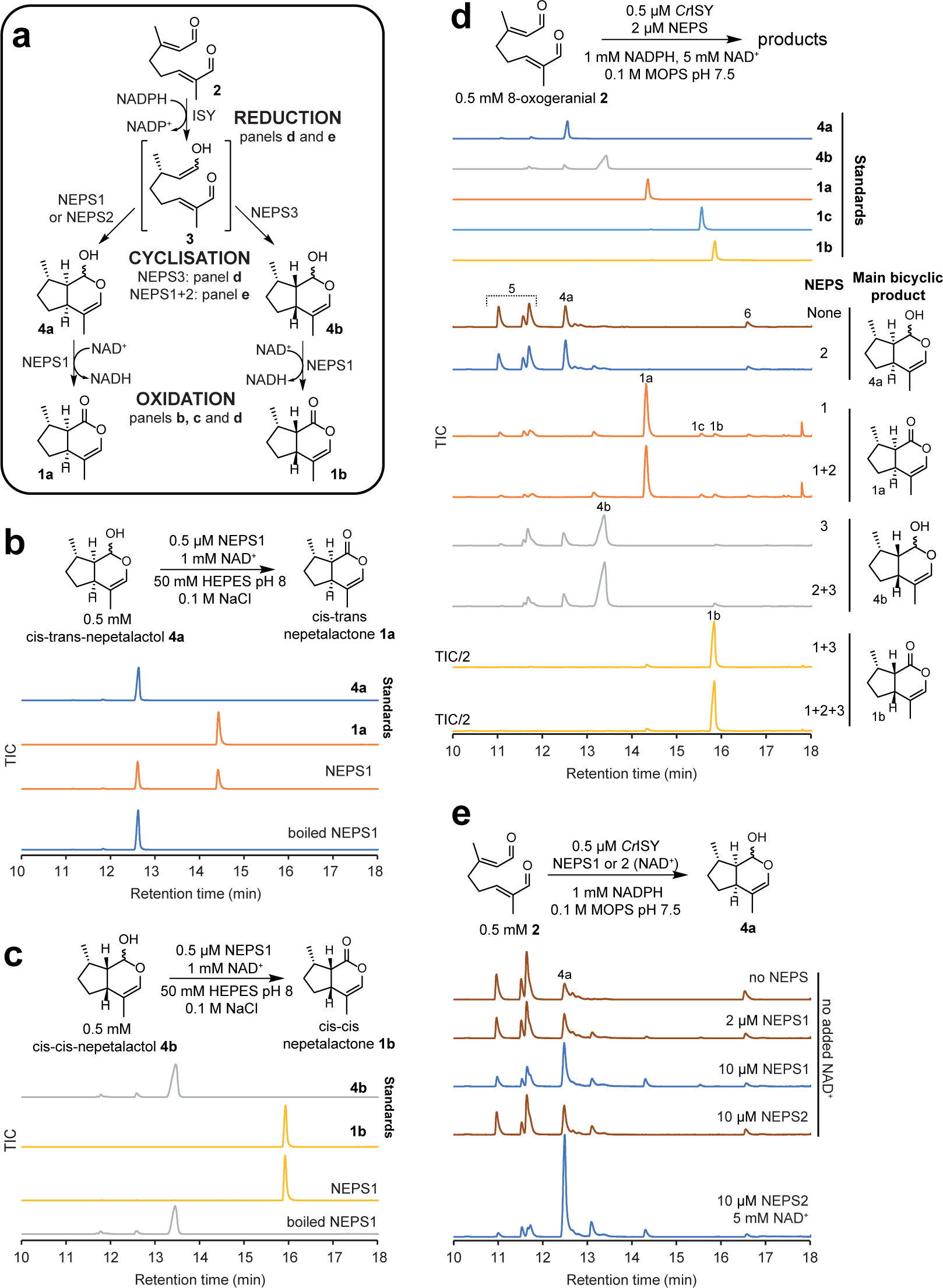
Formation of nepetalactones by NEPS enzymes. **a**, Summary of NEPS enzyme activities described in this figure. **b**, Cis-trans-nepetalactol dehydrogenase activity of NEPS1. NEPS1 catalyses the NAD^+^-dependent dehydrogenation of cis-trans-nepetalactol **4a** to cis-trans-nepetalactone **1a**. **c**, Cis-cis-nepetalactol dehydrogenase activity of NEPS1. NEPS1 catalyses the NAD^+^-dependent dehydrogenation of cis-cis-nepetalactol **4b** to cis-trans-nepetalactone **1b**. **d**, Combined activities of ISY and NEPS enzymes. Incubation of 8-oxogeranial **2**, *Cr*ISY, NEPS and cofactors enables the production of: cis-trans-nepetalactol **4a** (no NEPS or NEPS2), cis-trans-nepetalactone **1a** (NEPS1), cis-cis-nepetalactol **4b** (NEPS3) or cis-cis-nepetalactone **1b** (NEPS1 and NEPS3). See Supplementary Fig 11 for comparison of NEPS cascades with *Nm*ISY2 and *Cr*ISY. See Supplementary Fig 12 for further analysis of the NEPS3 cyclisation reaction. **e**, NEPS-catalysed formation of cis-trans-nepetalactol **4a**. Adjusting the NAD^+^ and/or NEPS concentrations reveals NEPS1 and NEPS2 can promote the formation of cis-trans-nepetalactol **4a**. All reactions were incubated for 3 h and are presented as GC-MS TICs. See Supplementary Fig 8 for EI spectra.

**Figure 4:**
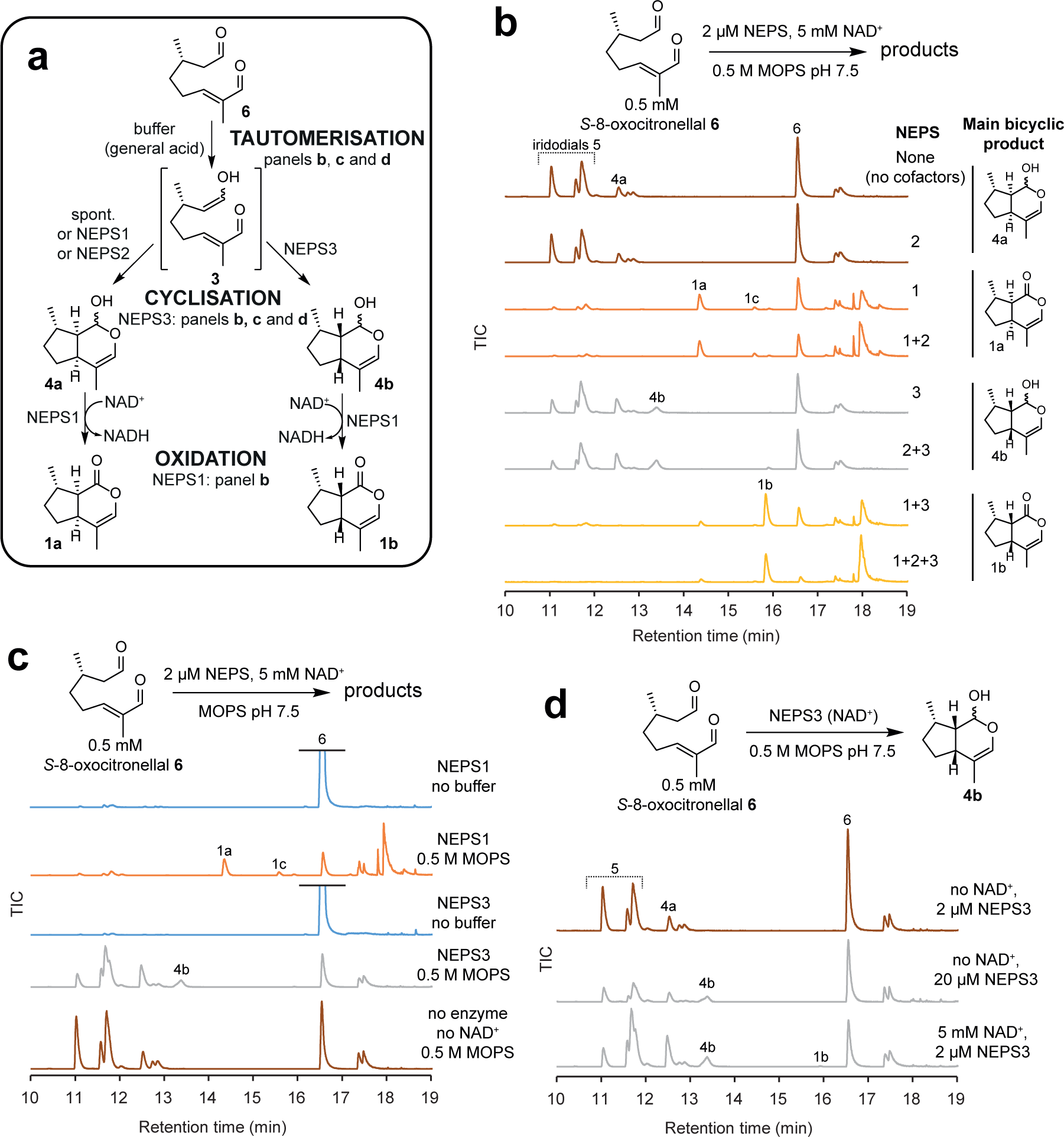
NEPS activities explored with *S*-8-oxocitronellal **6**. **a**, Summary of NEPS enzyme activities described in this figure. **b**, NEPS activities with *S*-8-oxocitronellal **6**, buffer and NAD^+^, presented as GC-MS TICs. The panel largely recapitulates observations of Figure 3b, but in the absence of ISY. Unknown side products are formed by NEPS1. See Supplementary Fig 13 for reactions with **6**, NEPS and *Cr*ISY. **c**, Buffer dependence of NEPS activity with *S*-8-oxocitronellal **6**. In the absence of buffer, NEPS1 and NEPS3 have no detectable activity; addition of buffer reveals enzyme activities. Buffer-catalysed tautomerisation of **6** appears to be necessary for enzyme activity, supporting the hypothesis that the activated 8-oxocitronellyl enol **3** and not *S*-8-oxo-citronellal **6** is the key NEPS substrate. **d**, NEPS3 catalysed cyclisation. The addition of NAD^+^ is not required for NEPS3 cyclisation activity, though addition does promote the reaction. We hypothesise that the cyclisation is not oxidoreductive, but NAD^+^ acts in a non-chemical manner (i.e. protein stabilisation). All reactions analyses are presented as GC-MS TICs.

The sensitivity of the product distribution to changes in solvent conditions led us to hypothesise that ISY reduces **2** to form the activated intermediate **3** which then leaves the enzyme active site and diffuses into the solvent. In the solvent, **3** can quench through either cyclisation or tautomerisation to form a mixture of products. We propose that the buffer acts as a general acid catalyst, promoting tautomerisation. Using MOPS buffer, this results in three regimes (Fig. 2c): at low buffer concentrations (<50 mM) two cyclisations generate the bicyclic **4a** as the dominant product; in moderately buffered concentrations (50-500 mM) one cyclisation followed by keto-enol tautomerisation to form monocyclic **5a** becomes favoured; at high buffer concentrations (≥500 mM) direct keto-enol tautomerisation of **3** into **6** becomes the dominant route. The promotion of tautomerisation by buffer molecules mirrors simulations that have highlighted the active involvement of solvent molecules in tautomerisation mechanisms^26,27^.

To validate that the ISY product profile was a result of the non-enzymatic cyclisation of **3**, we aimed to form **3** in the absence of enzyme. This was achieved by incubation of *S*-8-oxocitronellal **6** in unbuffered water at acidic (<2) or alkaline (>10) pHs (Supplementary Fig. 5), or in buffered water at pH 7.5 (Fig. 2d, Supplementary Fig. 6). Buffer or extreme pH promotes keto-enol tautomerisation of **6** into **3**, which could then undergo cyclisation/tautomerisation in an analogous manner to ISY reactions. In fact, formation of **4a** from **6** is an established synthetic route^28^. In high concentrations of buffer (500 mM MOPS), the product profiles of the ISY catalysed reduction of **2** and the non-chemical cyclisation of **6** are remarkably similar (Fig. 2d, Supplementary Fig. 6), supporting the hypothesis that iridoid cyclisation is not enzymatically catalysed.

**Figure 5:**
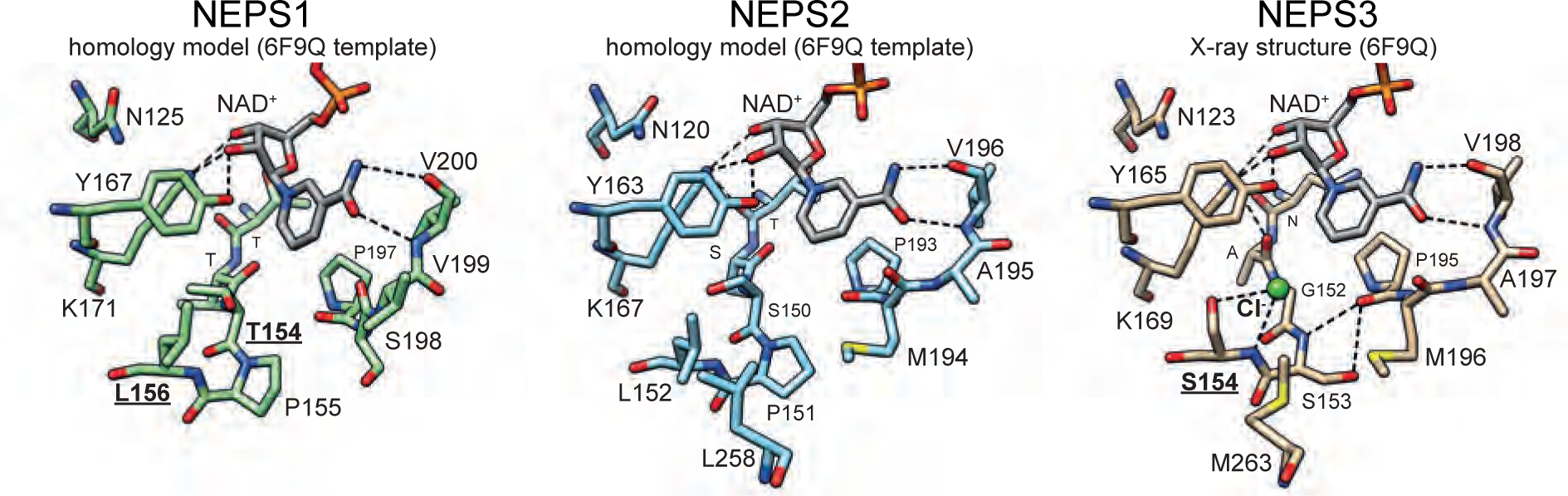
Structure of NEPS enzymes. X-ray crystal structure of NEPS3 (6F9Q) and homology model structures of NEPS1 and NEPS2. Active site NAD^+^ and residues are depicted as sticks. Dashed lines represent proposed H-bonds. The NEPS3 active site lacks the characteristic Ser/Thr of the SDR catalytic tetrad (G152). It also features a chloride bound to S154 and H-bonding between S153 and P190, features absent from NEPS1 (L156) and NEPS2 (L152). The role of residues in bold and underlined have been analysed by mutation. See Supplementary Fig 14 for further analysis of the NEPS3 crystal structure.

**Figure 6:**
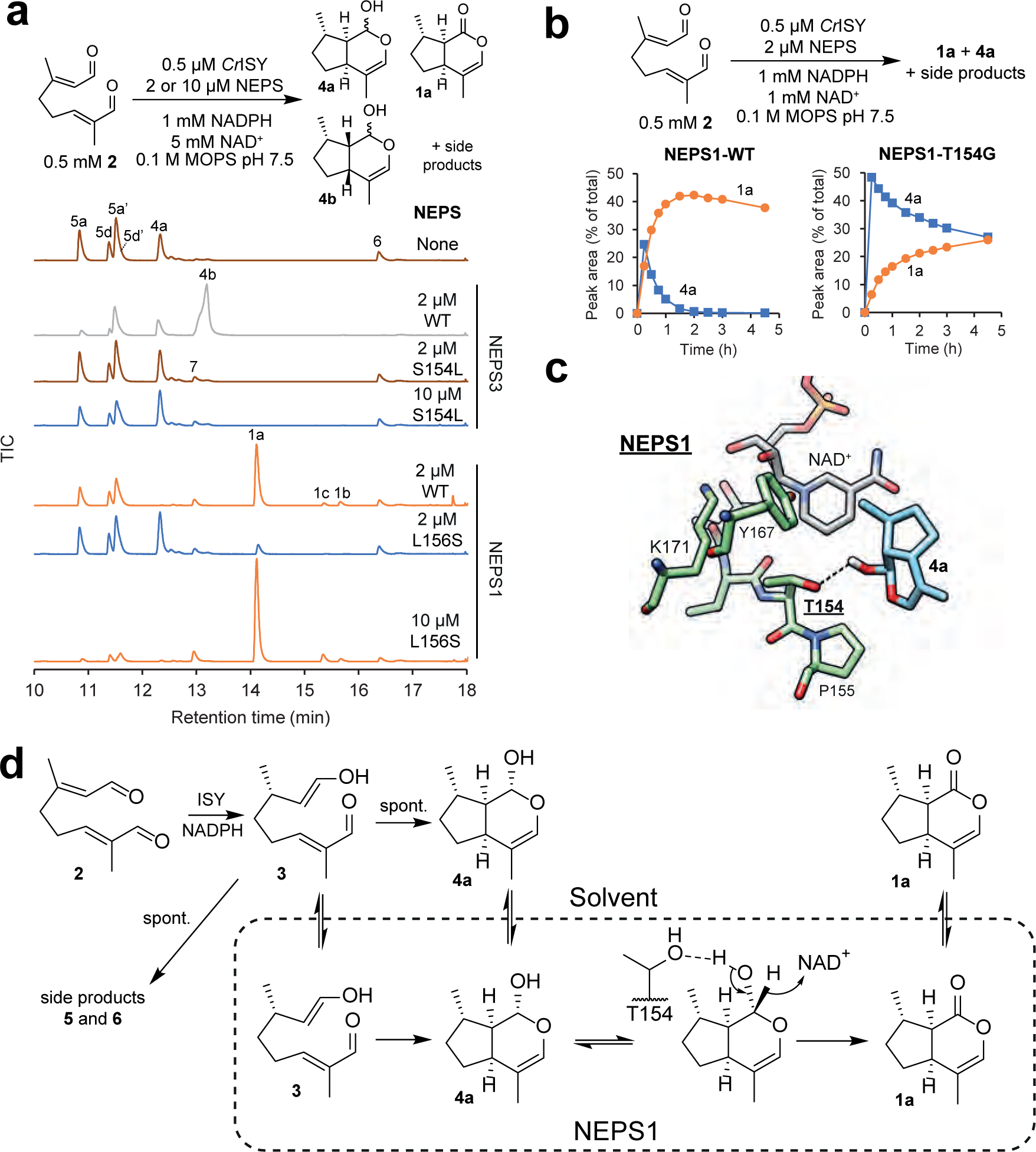
NEPS variants. **a**, Coupled assay with 8-oxogeranial **2**, ISY and NEPS variants, presented as GC-MS TICs. The product profile was measured after 3 h. The substitution S154L in NEPS3 appears to remove the cis-cis cyclase activity (absence of **4b**), but with 10 µM enzyme the formation of cis-trans-nepetalactol **4a** appears to be promoted relative to iridodials **5**. The L156S substitution in NEPS1 largely removes dehydrogenase activity of NEPS1 (less cis-trans-nepetalactone **1a**) but no increase in cis-cis-nepetalactol **4b** is observed. **b**, Time course of coupled assay with 8-oxogeranial **2**, ISY and NEPS1 WT or T154G. Quantities of cis-trans-nepetalactol **4a** (blue) and cis-trans-nepetalactone **1a** (orange) measured over the course of the reaction. Product proportions described as GC-MS TIC peak areas as a percent proportion of total product peak area. NEPS1-WT oxidises **4a** into **1a** rapidly whilst NEPS1-T154G oxidises **4a** with less efficiency. See Supplementary Fig 16 for all TICs of time course, and Supplementary Table 2 for kinetic analysis. **c**, Putative binding mode of cis-trans-nepetalactol **4a** in the NEPS1 active site. Binding mode generated by computational docking calculations using the NEPS1 homology model. The putative H-bond interaction between the lactol and T154 is highlighted with a dotted line. The depicted binding mode was ranked third of ten predicted binding modes (rank 1 score = −6.4 kcal.mol^-1^, depicted rank 3 energy = −6.3 kcal.mol^-1^). **d**, Scheme of NEPS1 activities and interactions. NEPS1 appears to have two distinct activities—cyclisation and dehydrogenation. The behaviour of the mutant T154G suggests that T154 is involved in dehydrogenation. The two activities appear distinct and may even involve different active site interactions. The slightly improved cyclase activity of T154G may be due to poor binding of **4a** which frees more enzyme for binding and cyclising **3** into **4a**.

Therefore, unlike canonical terpene synthases which catalyse concerted activation and cyclisation (Fig. 1b), ISY catalyses the activation of its linear substrate (8-oxogeranial **2** to 8-oxocitronellyl enol **3**) but does not appear to catalyse the subsequent cyclisation. Instead, we hypothesise that **3** diffuses out of the ISY active site into the solvent where it cyclises. The notion of free **3** raised the possibility that the iridoid stereochemistry may be defined by a partner cyclase enzyme, capable of accepting **3** as a substrate and catalysing diastereoselective cyclisation.

### Identification of nepetalactol related short-chain-reductases (NEPS)

Identifying a cyclase that works in partnership with ISY presents a challenge: since this proposed reaction is unprecedented, it is difficult to predict what type of enzyme family would catalyse such a cyclisation. However, nepetalactone biosynthesis in *Nepeta* is localised to a specific plant organ, glandular trichomes^7^. Therefore, we could compare the proteome of trichomes to trichome-depleted leaves to identify genes that are selectively expressed at the site of nepetalactone biosynthesis, thereby considerably narrowing the pool of potential gene candidates.

We obtained proteomes for *Nepeta mussinii* trichomes, leaves and trichome-depleted leaves (Supplementary Fig. 7a, Supplementary Data). Comparison of these proteomes enabled identification of trichome enriched proteins. This approach was validated by the identification of trichome enriched enzymes from upstream isoprenoid biosynthesis (the 2-*C*-methylerythritol 4-phosphate (MEP) pathway) and iridoid biosynthesis (Supplementary Fig. 7b,c).

As a starting point, we initially used these proteomes to identify the enzyme that converts nepetalactol **4** to nepetalactone **1**, an NAD^+^ dependent enzyme. An enzyme with such activity had previously been isolated from the trichomes of *Nepeta mussinii* but its sequence was not identified^29^. Six trichome enriched dehydrogenase genes were cloned and recombinantly expressed in *E. coli* (Supplementary Fig. 7d, Supplementary Table 1). Of these, one demonstrated cis-trans-nepetalactol **4a** dehydrogenase activity (Supplementary Fig. 7e).

The active enzyme is a short-chain reductase (SDR), part of the SDR110C family, a large and diverse family of NAD-dependent dehydrogenases often associated with plant secondary metabolism^30^. Consequently, it was named <u>Nep</u>etalactol-related <u>S</u>DR 1 (NEPS1). NEPS1 could catalyse the NAD^+^-dependent dehydrogenation of either cis-trans-nepetalactol **4a** or cis-cis-nepetalactol **4b** to the corresponding nepetalactones **1a** and **1b** (Fig. 3a-c, Supplementary Fig. 8a, Supplementary Table 2). Whilst the overall catalytic efficiency was slightly greater for dehydrogenation of **4a**, **4b** was turned over with twice the rate. This ratio of activities was in accordance with characterisation of the native enzyme^29^.

Sequence analysis of NEPS1 and the remaining dehydrogenase candidates revealed two additional trichome enriched paralogs of NEPS1, NEPS2 and NEP3 (Supplementary Fig. 9a,b). Phylogenetic analysis revealed that these three proteins have a close evolutionary relationship and are found uniquely within the *Nepeta* lineage (Supplementary Fig. 9c). Therefore, we hypothesised that NEPS2 and NEP3 also play a role in nepetalactone biosynthesis. Consequently, NEPS1-3 enzymes were assayed with a variety of nepetalactone related compounds and precursors such as nepetalactol **4**, iridodials **5** and 8-oxogeranial **2** (Supplementary Fig. 10). However, besides the NEPS1 activities described above, no other notable activities were observed for NEPS1-3.

### NEPS activities in conjunction with ISY

As described above, mechanistic investigations of ISY led us to hypothesise that a separate cyclase enzyme may act on the activated intermediate **3** that is generated by ISY. To test whether NEPS act as such cyclases, we performed one-pot cascade reactions combining NEPS enzymes with the ISY catalysed reduction of 8-oxogeranial **2** (Fig. 3a, d, e). As anticipated, addition of NEPS1 and excess NAD^+^ led to the formation of cis-trans-nepetalactone **1a** (Fig. 3d, Supplementary Fig. 8b), though unexpectedly ISY iridodial side products **5** were diminished. Remarkably, addition of NEPS3 to ISY and 8-oxogeranial **2** led to the formation of cis-cis-nepetalactol **4b**. A combination of NEPS1 and NEPS3 led to the production of cis-cis-nepetalactone **1b**. Adjusting enzyme and cofactor concentrations revealed that NEPS1 and NEPS2 promoted the formation of cis-trans-nepetalactol **4a** at the expense of iridodial **5** (Fig. 3e). The products observed indicated that the NEPS enzymes are cyclases (Fig. 3a), capable of accepting **3**, the product of ISY, and cyclising it to **4a** (NEPS1 and 2) or **4b** (NEPS3). NEPS1 can then oxidise **4a** and **4b** into **1a** and **1b** respectively. The activities of NEPS enzymes did not appear to differ when tested with *Cr*ISY or *Nm*ISY2, suggesting that protein-protein interactions between ISYs and NEPS do not play a role in the system (Supplementary Fig. 11).

The cyclisation of **3** into **4** is a non-oxidoreductive net [4+2] cycloaddition. The cyclase activities of NEPS also do not appear to be oxidoreductive. Investigation into the co-factor dependence of the ISY-NEPS3 reactions demonstrated this: NAD^+^ concentrations were not limiting to overall reaction conversions, suggesting that NAD^+^ was not consumed (Supplementary Fig. 12). Furthermore, NEPS3 was active in the absence of supplemented NAD^+^, though addition did improve activity. It appeared that although NAD^+^ is not turned over by NEPS3 during cyclisation it may promote the enzyme’s catalytic ability, perhaps through stabilisation of the protein structure.

### NEPS activities with *S*-8-oxocitronellal

To verify the NEPS activities, reactions were conducted with *S*-8-oxocitronellal **6** and without ISY (Fig. 4). High concentrations of buffer were employed to promote the formation of **3**, the proposed NEPS substrate, from **6**. In these conditions, the previously observed activities of NEPS were recapitulated (compare Fig. 4b to Fig. 3d).

Neither NEPS1 nor NEPS3 were active when incubated with **6** in the absence of buffer (Fig. 4c). Buffer was necessary for activity, supporting the notion that that **6** is not the key substrate, but the tautomer **3** is. Further evidence for this was obtained by adding *Cr*ISY to reactions containing NEPS and **6**—the pattern of products observed implied NEPS were binding to **6** without turning it over (Supplementary Fig. 13).

Support for the non-oxidoreductive nature of the NEPS3 cyclisation came from incubating NEPS3 with **6** and buffer in the absence of supplemented NAD^+^ or NADPH: formation of cis-cis-nepetalactol **4b** was still observed (Fig. 4d). As noted above, addition of NAD^+^ does promote the reaction, though it is not necessary for activity. Interestingly, at high NAD^+^ concentrations, trace quantities of **1b** are observed (Fig. 4d, also Fig. 3d). Overall, NEPS reactions with **6** support the notion of **3** as the substrate and add further evidence to a non-oxidoreductive cyclisation (Fig. 4a).

### Structure and mechanism of NEPS enzymes

To understand the mechanism of the NEPS3 cis-cis cyclisation reaction, we obtained an X-ray crystal structure of NEPS3 bound to NAD^+^ (6F9Q, Supplementary Table 3). In common with structural homologs, it forms a homotetramer with 222 symmetry, with the four active centres contained entirely within individual protomers (Supplementary Fig. 14a). Efforts to generate an *apo* or ligand bound structure were unsuccessful, as were efforts to crystallise NEPS1. Despite the fact that NEPS3 appears to function primarily as a non-oxidoreductive cyclase, its structure is characteristic of classical SDRs, with NAD^+^ bound in the typical fashion (Supplementary Fig. 14b,c). Although evidence suggests NAD^+^ is not turned over by NEPS3 during cyclisation, the co-factor appears important for the enzyme structure as it was required for crystallisation and it is bound by multiple H-bonds by the protein.

Homology models of NEPS1 and NEPS2, based on the NEPS3 structure, were generated to enable comparison of the enzyme active sites (Fig. 5). Whilst NEPS1 and NEPS2 have a typical SDR catalytic tetrad (N-Y-K-T/S), NEPS3 lacks the theronine/serine and instead has a glycine (G152). In addition, NEPS3 features H-bonding between active site residues S153 and P190, absent from both NEPS1 and NEPS2 due to the presence of a proline in place of the serine.

Interestingly, the NEPS3 structure appeared to feature a chloride anion bound to the amide NH and side chain of S154. Based on the presence of the chloride, and the similarity to the substrate oxyanion binding site in *Cr*ISY, this position may be a substrate oxyanion binding site (Supplementary Fig. 14c). This site appears to be absent in NEPS1 and 2 due to the steric hindrance of the leucine side chain present in the equivalent position.

Active site residue roles in NEPS1 and NEPS3 were probed via mutational screens (Supplementary Table 4). NEPS3 was especially sensitive to mutation, with several active site substitutions abolishing detectable soluble expression (A151T, G152T, S153P, Y165F). These mutations may have disrupted the H-bonding network in the active site, reducing protein stability. Of the mutations yielding soluble protein, only N150T maintained native levels of activity, whilst S154L, K169M and M196S had a severe reduction or complete loss of activity (Supplementary Fig. 15a). Several NEPS1 variants had no detectable formation of **1a**, signifying that these may be functionally important residues (T152N, L156S, Y167F, K171M, S198M, T202A, Supplementary Fig. 15b). Near-WT levels of **1a** were identified in samples with the variants V110A, T153A, V199A, whilst trace quantities of **1a** was identified with N125A and P155S (Supplementary Fig. 15c). Most interesting were variants containing the T154G substitution, which produced considerably greater quantities of **4a** than WT (Supplementary Fig. 15d).

The role of the NEPS3 S154 putative oxyanion binding site was examined by further characterisation of NEPS3-S154L and the complementary NEPS1-L156S variant. NEPS3-S154L (2 µM) demonstrated no detectable cis-cis cyclase activity, though at higher enzyme concentrations (10 µM), the variant appeared to promote the formation of cis-trans-nepetalactol **4a** (Fig. 6a). The complementary substitution in NEPS1, L165S, reduced dehydrogenase activity but failed to increase the formation of cis-cis-nepetalactol **4b** or lactone **1b** (Fig. 6a). The NEPS3-S154 oxyanion binding site therefore appears necessary for cis-cis cyclisation in NEPS3 but its introduction into NEPS1 is not sufficient to establish cis-cis cyclisation activity. In contrast, the removal of S154 from NEPS3 appears to switch the cyclisation selectivity from cis-cis to cis-trans.

The NEPS1-T154G variant was also characterised further. Compared to NEPS1-WT, the variant accumulated **4a** and showed impaired formation of **1a** (Fig. 6b, Supplementary Fig. 16a). This loss of dehydrogenase activity appears to be a result of a 150-fold increase in K_m_ for **4a** caused by the T154G substitution (Supplementary Table 2). This result indicates that T154 interacts with **4a** during or prior to the dehydrogenation step; a putative binding mode is provided by docking calculations (Fig. 6c). Interestingly, the variant appears to have slightly enhanced cyclisation activity: reactions with T154G show minimal formation of cis-trans-iridodial **5a** side products (Supplementary Fig. 16a) and the variant is also able to form detectable levels of **4a** from **6** in buffer without supplemented cofactors (Supplementary Fig. 16b). This improved cyclisation may be a consequence of poor **4a** binding—if **4a** is released rapidly then there is more available enzyme to bind **3** and catalyse its cyclisation (Fig. 6d).

## Discussion

Here we demonstrate how a mechanistic analysis of ISY led to the hypothesis that a separate cyclase is responsible for setting the stereochemistry of the iridoid framework. A comparative proteomic analysis allowed discovery of these cryptic enzymes. These NEPS enzymes demonstrate the plasticity and innovation characteristic of plant secondary metabolism enzymes^31^. NEPS3, for example, is an SDR by structure and sequence yet its primary catalytic activity is non-oxidoreductive; NAD^+^ is utilised not as a co-substrate but as a protein structural scaffold. Interestingly, at very high NAD^+^ concentrations NEPS3 can catalyse dehydrogenation with low catalytic efficiency, a phenomenon that may be a glimpse into its evolutionary past.

Due to its inherent reactivity, *in situ* generation of the substrate **3** was required to reveal the activity of the NEPS enzymes. Reactive non-isolable substrates appear elsewhere in plant specialised metabolism, including monoterpene indole alkaloid^32^ and lignan biosynthesis^33,34^. NEPS are reminiscent of dirigent proteins, proteins in lignan biosynthesis that control the stereoselective cyclisation of a reactive intermediate that is generated by a separate enzyme^33,34^. There may be similar undiscovered steps in other metabolic pathways; these cannot be revealed in simple one substrate, one enzyme assays, but require multi-enzyme cascade strategies, making discovery of such enzymes challenging.

Based on the structural and mutant data, we propose that NEPS1/2 and NEPS3 catalyse cyclisation in different fashions. NEPS1/2 appear to allow the enol **3** to proceed down a default ‘uncatalysed’ path, forming the same product as formed in water (Fig. 2). Their key role in the cyclisation appears to be to protect the intermediate from general acid catalysed tautomerisation; there is no evidence to suggest that NEPS1/2 mechanisms do not mirror the stepwise Michael addition seen in solution^35^. NEPS3, on the other hand, has specific 7*S*-cis-cis-nepetalactol cyclase activity, binding to **3** possibly via S154 and exerting steric and/or electrostatic influence to enable formation of the cis-cis stereochemistry. Further analysis is required to determine whether NEPS3 catalyses the [4+2] cyclisation in a stepwise or concerted manner (i.e. Michaelase or Diels-Alderase). The contrast between the ‘passive’ NEPS1/2 and ‘active’ NEPS3 mechanisms mirrors the of dichotomy of intrinsic substrate reactivity versus enzyme influence in canonical terpene synthase mechanisms^13^.

Phylogenetic analysis suggests the NEPS enzymes are unique to *Nepeta*. Yet ISYs from other organisms also do not catalyse iridoid cyclisation. Thus, it is likely that different cyclases, unrelated to NEPS, are operating in other iridoid producing species. The absence of such cyclases from iridoid pathways reconstituted in microbial organisms may have negatively impacted yields^36–38^. The NEPS cyclases described here, especially NEPS2, a dedicated cis-trans cyclase, can be incorporated into such systems to improve yield.

We have revealed the biosynthetic origin of cis-trans and cis-cis-nepetalactone, compounds in *Nepeta* responsible for cat attraction and insect repellence. In doing so we have discovered three novel enzymes: two dedicated cyclases and one multifunctional cyclase-dehydrogenase. The structure of one of these enzymes reveals it has re-purposed a dehydrogenase structure for a different catalytic function. We have shown that iridoid biosynthesis involves uncoupled activation and cyclisation: a reactive, non-isolable enol is formed by reduction and cyclised by separate enzymes. Such findings will contribute to synthetic biology metabolic reconstructions and inform the *de novo* design of (bio)synthetic pathways; they also highlight the dynamic and innovative nature of plant natural product biosynthesis.

## Data availability

The sequences of *N. mussinii* NEPS enzyme have been deposited in GenBank/EMBL/DDBJ with the accession codes: MG677124 (NmNEPS1), MG677125 (NmNEPS2) and MG677126 (NmNEPS3). The NAD^+^ bound NmNEPS3 (7*S*-cis-cis-nepetalactol cyclase) X-ray structure has been deposited in the PDB with the accession code 6F9Q. The mass spectrometry proteomics data have been deposited to the ProteomeXchange Consortium via the PRIDE^39^ partner repository with the dataset identifier PXD008704. Detailed experimental procedures and can be found in the supplementary information. The authors declare that all other data supporting the findings of this study are available within this Article and its Supplementary Information or from the authors upon reasonable request.

## Acknowledgements

We acknowledge funding from UK Biotechnological and Biological Sciences Research Council (BBSRC) and Engineering and Physical Sciences Research Council (EPSRC) joint-funded OpenPlant Synthetic Biology Research Centre (BB/L014130/1) and from the National Science Foundation Plant Genome Research Program (IOS-1444499). For the X-ray data collection, we acknowledge Diamond Light Source for access to beamline I03 under proposal MX13467, with support from the European Community’s Seventh Framework Program (FP7/2007– 2013) under grant agreement 283570 (BioStruct-X). We are grateful to: Paul Brett (John Innes Centre) for assistance with GC-MS analysis, Marielle Vigoroux (John Innes Centre) for assistance with proteome annotations, and Nathaniel Sherden and Hajo Kries for providing chemical and genetic material. We also thank Kendall Houk, Jason Fell, Hajo Kries and Daniel Whitaker for discussions concerning the iridoid synthase and cyclisation mechanisms.

## Author Contributions

S.E.O’C. designed and supervised the project; B.R.L. performed molecular cloning, protein purification, enzyme assays, trichome isolation, chemical synthesis, phylogenetic analysis, homology modelling and computational docking; G.S. performed proteome analysis; M.O.K. assisted with protein purification, compound isolation and chemical synthesis; B.R.L., G.R.T. and C.E.M.S. performed crystallisation trials and obtained crystals; B.R.L., G.R.T. and D.M.L. refined structures; B.R.L and S.E.O’C. wrote the manuscript.

